# RUDEUS, a machine learning classification system to study DNA-Binding proteins

**DOI:** 10.1101/2024.02.19.580825

**Authors:** David Medina-Ortiz, Gabriel Cabas-Mora, Iván Moya-Barría, Nicole Soto-Garcia, Roberto Uribe-Paredes

**Affiliations:** Departamento de Ingeniería En Computación, Universidad de Magallanes, Avenida Bulnes 01855, Punta Arenas, Chile; Centre for Biotechnology and Bioengineering, CeBiB, Universidad de Chile, Beauchef 851, Santiago, Chile; Departamento de Química, Universidad de Magallanes, Avenida Bulnes 01855, Punta Arenas, Chile

**Keywords:** DNA-binding proteins, single-stranded and double-stranded DNA, Machine learning, Protein language models

## Abstract

DNA-binding proteins are essential in different biological processes, including DNA replication, transcription, packaging, and chromatin remodelling. Exploring their characteristics and functions has become relevant in diverse scientific domains. Computational biology and bioinformatics have assisted in studying DNA-binding proteins, complementing traditional molecular biology methods. While recent advances in machine learning have enabled the integration of predictive systems with bioinformatic approaches, there still needs to be generalizable pipelines for identifying unknown proteins as DNA-binding and assessing the specific type of DNA strand they recognize. In this work, we introduce RUDEUS, a Python library featuring hierarchical classification models designed to identify DNA-binding proteins and assess the specific interaction type, whether single-stranded or double-stranded. RUDEUS has a versatile pipeline capable of training predictive models, synergizing protein language models with supervised learning algorithms, and integrating Bayesian optimization strategies. The trained models have high performance, achieving a precision rate of 95% for DNA-binding identification and 89% for discerning between single-stranded and doublestranded interactions. RUDEUS includes an exploration tool for evaluating unknown protein sequences, annotating them as DNA-binding, and determining the type of DNA strand they recognize. Moreover, a structural bioinformatic pipeline has been integrated into RUDEUS for validating the identified DNA strand through DNA-protein molecular docking. These comprehensive strategies and straightforward implementation demonstrate comparable performance to high-end models and enhance usability for integration into protein engineering pipelines.

## Introduction

The interactions between DNA and proteins form the basis of numerous cellular processes crucial for biological functions (Zhang et al., 2022). Approximately 6-7% of eukaryotic proteins are known to interact with DNA. These proteins possess distinctive DNA-binding domains and different affinities for single-stranded and double-stranded DNA (Attali et al., 2021; Gupta et al., 2021). The DNA-protein recognition mechanisms involve direct base–amino acid interactions and indirect contributions from conformational energy derived from DNA deformations and elasticity (Arora et al., 2023).

DNA-binding proteins play relevant roles in diverse biological processes, including DNA replication, transcription, packaging, and chromatin remodelling (Kabir et al., 2024). These proteins guide strand separation, maintain DNA integrity, regulate gene expression, compact genetic material, and reorganize chromatin structure. Understanding the characteristics and functions of DNA-binding proteins has become essential across various scientific disciplines (Zhang et al., 2022). Identifying these proteins enhances its comprehension of structural regulations and facilitates understanding of relations between gene mutations and genetic diseases (Kabir et al., 2024; Alendar and Berns, 2021a). The exploration of DNA-binding proteins, such as TDP-43, helicase chromodomain proteins, and those with methyl-CpG-binding domains, has notably contributed to recognizing pathologies like neurodegenerative disorders and cancers (Lye and Chen, 2022; Alendar and Berns, 2021b; Wang et al., 2022b).

Advances in computational biology and bioinformatics have significantly facilitated the study of protein functionality, allowing for comprehensive analyses of behaviour, structure, interactions, and physicochemical properties (Fu et al., 2020). While these computational tools have greatly supported molecular biology research, their limitations in conducting large-scale functional studies have required the integration of artificial intelligence (AI) strategies and machine learning algorithms (ML) (Wang et al., 2022c). This integration has accelerated the discovery of new DNA-binding proteins, focusing on predicting interaction sites, hotspots, and transcription factor binding sites (Wang et al., 2022a; Pan et al., 2020; Zhang et al., 2021b).

Various approaches have been employed for DNA-binding protein recognition using ML methods, ranging from classic strategies with feature engineering to more recent implementations of deep learning architectures (Shadab et al., 2020; Zhang et al., 2020; Ali et al., 2022; Banjar et al., 2022; Barukab et al., 2022). Despite achieving similar performances, comparing methods is challenging due to variations in training datasets and validation examples. In the context of classifying single-stranded and double-stranded DNA, similar methodologies have been applied to develop predictive systems (Wang et al., 2017; Ali et al., 2020; Tan et al., 2019) generating similar challenges, with limited exploration of physicochemical encoders and numerical representation strategies supported by large language models (Medina-Ortiz et al., 2022; Medina et al., 2023; Fernández et al., 2023).

This paper introduces RUDEUS, a Python library for DNA-binding classification systems and recognising single-stranded and double-stranded interactions. RUDEUS incorporates a generalizable pipeline that combines protein language models, supervised learning algorithms, and hyperparameter tuning guided by Bayesian approaches to train predictive models. Using this pipeline, it was trained and validated two classification models for DNA-binding identification and single-stranded and double-stranded interaction evaluation, achieving precision rates of 95% and 89%, respectively. RUDEUS’s usability is demonstrated by evaluating various DNA-binding proteins, annotating over 20,000 protein sequences, and validating them using structural bioinformatic approaches. The developed strategies and RUDEUS’s implementation and simplicity showcase comparable performances to high-performance models and offer enhanced usability for latent space exploration and mutational landscape navigation in DNA-binding proteins.

## 2 Methods

### 2.1 Collecting and processing protein sequences

All protein sequences were collected from the literature, databases, and previously reported works. The datasets reported in (Hu et al., 2019; Shadab et al., 2020) were employed for the case of DNA-binding protein classification models. In contrast, the dataset reported in (Sharma et al., 2021; Wang et al., 2017) was utilized for the single-stranded and double-stranded DNA-binding protein classification models. Once the datasets were collected, a Python script was implemented to join and filter redundancy. Finally, a length filter was applied to remove sequences with more than 1024 residues and less than 50 residues and remove sequences with non-canonical residues. This filter is only to facilitate the application of the different pre-trained models to numerical representation strategies. In summary, two datasets were constructed for DNA-binding protein classification models and for single-stranded and double-stranded DNA-binding protein classification models, with 47500 (26982 negative and 20518 positive examples) and 1233 examples (965 double-stranded and 268 single-stranded), respectively.

### 2.2 Numerical representation strategies

Once the datasets are processed and compiled, numerical representation strategies are applied to obtain numerical vectors for training the predictive models. This work employs different pre-trained models based on protein language models, including prottrans family model (Elnaggar et al., 2020) and ESM family models (Rives et al., 2021; Meier et al., 2021). All pre-trained models were applied through the bio-embedding tool, combined with a reduction process to obtain vectors in a 1 *− D* dimension (Dallago et al., 2021).

### 2.3 Training predictive models and tuning optimization

A classic machine learning pipeline was employed to train predictive models (Medina-Ortiz et al., 2020). First, the dataset is divided into training and validation datasets using a 70:30 proportion. Then, a supervised learning algorithm is used with default hyperparameters to train a predictive model employing the training dataset. The validation dataset is utilized to obtain the performances of the training process. RUDEUS has different supervised learning algorithms, including random forest, decision trees, support vector machine, KNN, AdaBoost, ExtraTree, Gaussian Process, XGBoost, and Gradient Boosting. Besides, a *k* −fold crossvalidation *k* = 10 was employed to prevent overfitting, estimating a performance metric during the training and validation. Finally, a fine-tuning process is generated to recognize the better hyperparameters to optimize the performance metrics. This process is based on a Bayesian approach and implemented through the Optuna Library (Akiba et al., 2019). RUDEUS exports the models as a joblib instance and generates a summary report of the training process, including dataset details, performances, and time process. Alternatively, a balanced process is incorporated for an unbalanced dataset. This process creates a balanced dataset by applying undersampling strategies.

### 2.4 Using predictive models and evaluating predictions through structural bioinformatic approaches

The classification models implemented in RUDEUS are available to use stand-alone or employ the modules available in RUDEUS. In both cases, protein sequences are necessary as input to work; then, the numerical representation strategy needs to be applied to obtain numerical vectors. Once the protein sequences are coded, the model is loaded, and predictions can be generated. The model gives predictions depending on the type of response. For example, if it is used the DNA-binding classification system, the predictive model could generate two types of responses: positive (the protein is DNA-binding) and negative (the protein is not DNA-binding). Besides, you can use the single-stranded and double-stranded DNA classification with DNA-binding proteins to identify the type of union with DNA molecules. In this case, it is necessary to do the same steps: load the model, apply the numerical representations, and predict with the loaded model. It highly recommends hierarchically using the models, first evaluating whether a protein is DNA-binding and then evaluating the type of binding.

Moreover, a structural bioinformatic pipeline is incorporated in RUDEUS to evaluate the generated predictions by the model using a DNA-protein molecular docking through the LightDock v9.4 software (Roel-Touris et al., 2020). This pipeline requires the protein and DNA structures in PDB format as input. First, the implemented pipeline prepares the structures applying: i) a protonation for each input, ii) elimination of previous hydrogens available in the structures, iii) rebuilding the structure using the reduce library (Roel-Touris et al., 2020), and iv) a modify names or remove incompatibles atoms the AMBER94 force field, followed by an update in the atoms numeration. Once the preparation structures are done, the molecular docking is started with initial configurations: i) 400 swarms, ii) 200 glowworms, and iii) 100 steps for performing docking calculations. Then, the generated conformers are clustered based on the RMSD metric employing the BSAS function. Finally, the best pose is selected based on the highest docking score.

### 2.5 Availability and Implementation strategies

All source code was implemented under the Python Language programming v3.9.16, including the modules, libraries, and demonstration scripts in RUDEUS. The main libraries employed to develop the predictive models were scikit-learn (Pedregosa et al., 2011) and Optuna (Akiba et al., 2019). Furthermore, to process and compile all datasets, the Pandas library was employed (McKinney et al., 2011). Finally, a conda environment was constructed to facilitate the deployment of the built library, combined with different Jupyter Notebooks, to ensure the replicability of the presented work. All source code, environment configuration, datasets, and created models are available for noncommercial uses in the GitHub repository https://github.com/ProteinEngineering-PESB2/DNA-Binding-modelshttps://github.com/ProteinEngineering-PESB2/DNA-Binding-models under the MIT licence.

## 3 Results and discussions

### 3.1 RUDEUS trains its predictive models using a generalizable machine learning pipeline

RUDEUS is a Python library that facilitates the implementation of predictive models for DNA-Binding protein identification, discovery, and evaluation through bioinformatic approaches. All predictive models available in RUDEUS are trained and validated using the same machine-learning pipeline. Figure 1 summarizes the implemented pipeline and the main steps.

**Figure 1:**
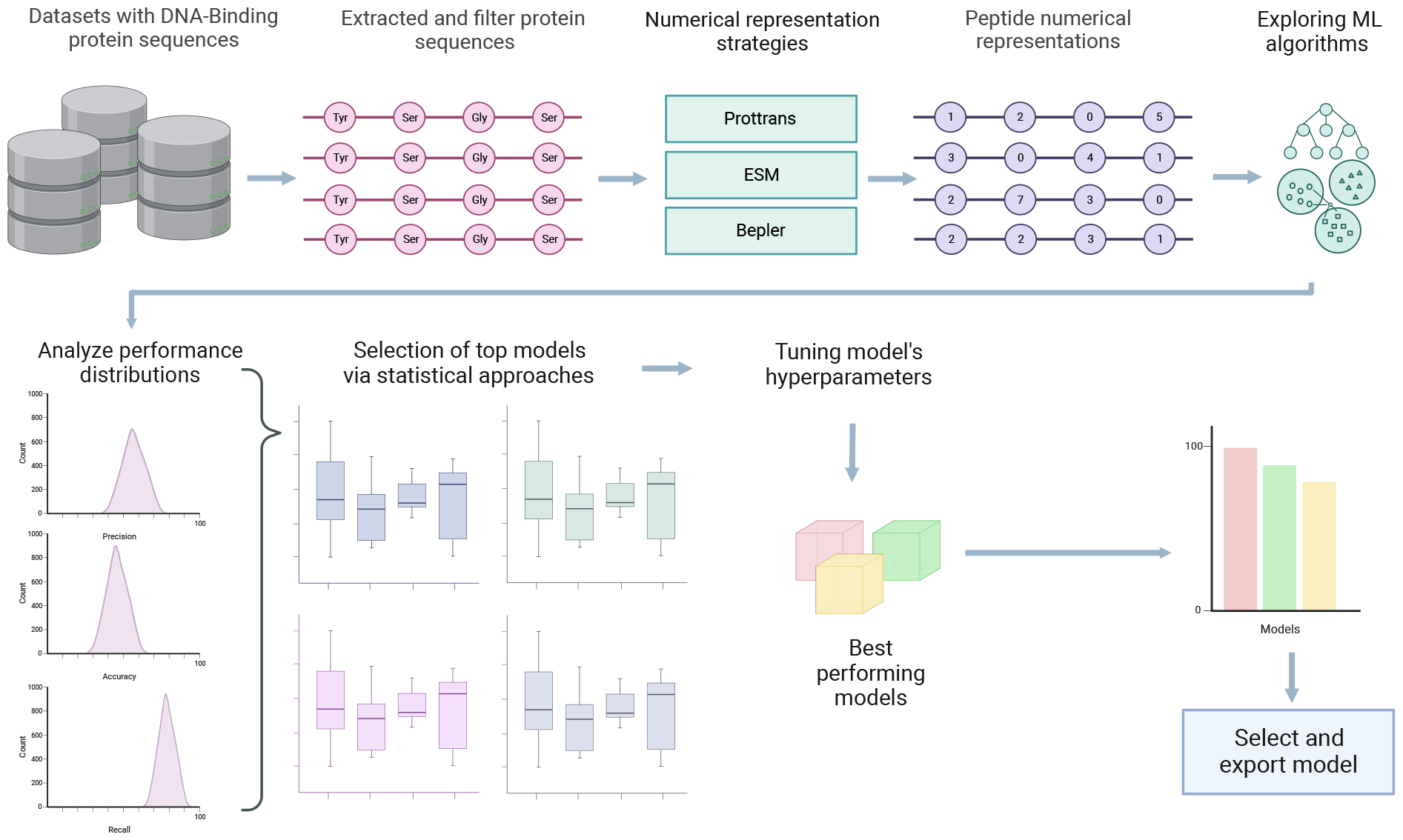
The designed e implemented pipeline to train predictive models for DNA-Binding identification incorporated in RUDEUS. The proposed pipeline first collects and processes the protein sequences by incorporating length filters and removing non-canonical residues. Then, numerical representation strategies are applied to obtain encoded vectors through pre-trained models based on protein language models, including Prottrans family models, ESM family models, Bepler, Glove, and all the different pretrained models available in the bio-embedding library. Then, different supervised learning algorithms are explored using default hyperparameters employing all generated datasets in the previous step. Then, statistical approaches are applied to filter and select the best combinations of supervised learning algorithms and numerical representation approaches. A Bayesian approach guides the selected combinations tuning hyperparameters process through the Optuna library, and ensemble learning is explored to evaluate different combinations of the individual optimized models. Finally, the best strategy is selected based on the best performances, including training, validation, and overfitting ratio.

First, the DNA-binding proteins are collected and processed applying filters: i) a length filter (protein sequences with a length higher than 50 residues and lower than 1024 residues) and ii) removing non-canonical residues. Once the filters are applied, the protein sequences are represented numerically using 16 pre-trained models available in the bio-embedding library (Dallago et al., 2021), including, for example, prottrans family models, Bepler, ESM family models, and Word2Vec. RUDEUS doesn’t incorporate amino acid coding approaches like physicochemical encoding and Fourier Transform to prevent the application of *zero-padding* techniques (Medina-Ortiz et al., 2022).

With the protein sequences numerically represented, the next step is the training process. As was previously described in the methodology section, a classic pipeline to train predictive models is applied (Medina-Ortiz et al., 2020) with the exception that the division of training and validation datasets is generated *n* = 100 times with different seeds to obtain metric distributions for each metric applied to evaluate the performances of the combinations of numerical representation approaches and supervised learning algorithm. Besides, if the dataset is unbalanced, an under-sample strategy is incorporated to generate a balanced dataset and then apply the training module.

Once the models are trained and the performance metrics are generated, a statistical approach is applied to filter and select the best combination strategies (supervised learning algorithm and numerical representation approach). The statistical selector includes: i) Get statistical values (mean and standard deviation) from the performance distribution for each applied metric in training and validation steps. ii) Define a Bernoulli event as a performance value on average higher than a specific quantile value. iii) Define a second Bernoulli event as a standard deviation value lower than a specific quantile value and generate a binomial distribution for all Bernoulli events. In total, 16 Bernoulli events are generated, combining average, standard deviation, and the four performance metrics employed to evaluate the model (precision, recall, accuracy, and F-score). iv) Finally, obtain the number of successes, generate the distribution of success, and obtain the positive outliers.

The designed statistical filter facilitates the identification of robust predictive models due to the low standard deviation values in their performance distribution and models with high performances due to the high average values in their performance distribution.

The selected models are optimized through a tuning optimization strategy. RUDEUS incorporates a Bayesian approach for tuning hyperparameters employing the Optuna library (Akiba et al., 2019). Different grids are generated depending on the supervised learning algorithm. Besides, the target optimization could be parametrized. However, this work uses the recall metric as an optimization target.

At last, the final strategy is selected by applying the following criteria: i) highest performances, ii) lowest overfitting, and iii) simplicity. The last criterion implies that if two models are similar in performance and overfitting rate, the most simple and easy to explain will be selected.

### 3.2 RUDEUS achieves high performances in its classification models

Two classification tasks were explored and evaluated in RUDES: DNA-binding protein classifications and single-stranded or double-stranded DNA type interaction. More than 10000 combinations of numerical representation approaches and supervised learning algorithms were generated for each task. Four metrics were employed to evaluate the performances of the predictive models, including accuracy, precision, recall, and F-score.

Figure 2 shows the distributions obtained for the exploration steps in the training process for recall metric. On average, the performances are 83% of precision for DNA-binding protein classification and 82% of precision for single-stranded or double-stranded DNA type interaction. The highest distribution performances in the DNA-binding classification model are achieved by applying the pro-trans Uniref, BDF, and XLU50 pre-trained models, independent of the supervised learning algorithm. In the case of single-stranded or double-stranded DNA type interaction models, the highest distribution performances are achieved by the pre-trained models pro-trans XLU50, Uniref, t5bdf, ESM1B, and ESM1V pre-trained models. Concerning the supervised learning algorithms, in both models, the highest distribution performances are achieved by ensemble algorithms like Random Forest, Gradient Boosting, ExtraTrees, and KNN methods.

**Figure 2:**
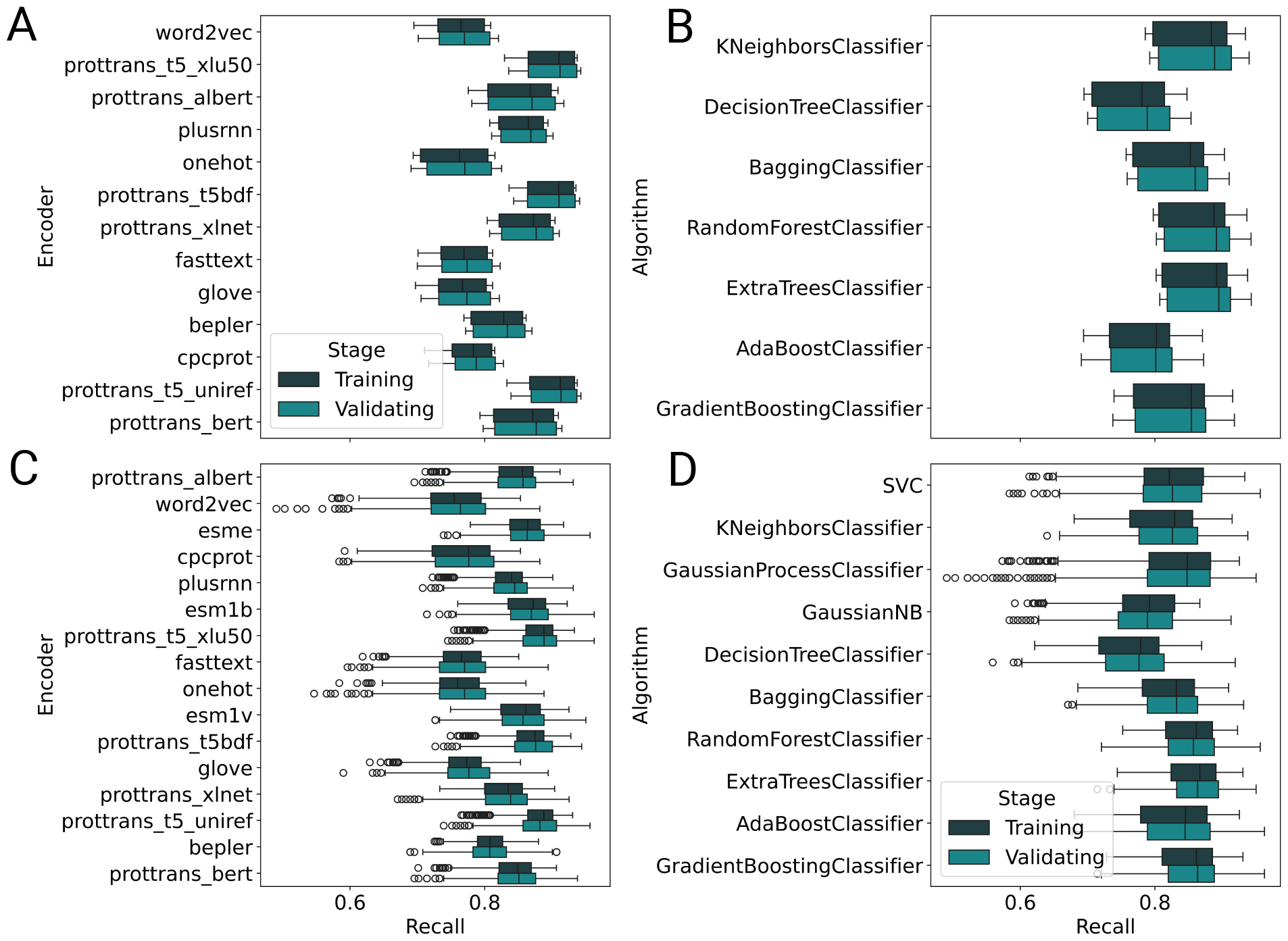
Recall distribution performances for all explored tasks in this work evaluated by numerical representation strategies and supervised learning algorithms. **A** Recall distribution for DNA-binding classification task grouped by pre-trained model employed as numerical representation strategy. **B** Recall distribution for DNA-binding classification task grouped by supervised learning algorithm. **C** Recall distribution for single-stranded or double-stranded DNA type interaction task grouped by pre-trained model employed as numerical representation strategy. **D** Recall distribution for single-stranded or double-stranded DNA type interaction task grouped by supervised learning algorithm.

The statistical selection process designed in this work was applied to obtain the best combinations of numerical representation strategies and supervised learning algorithms. In both cases, the sixteen Beronulli events were evaluated using a filter of the highest performances of all values over 90 quantiles general distribution and a filter of the robust model of all distributions with standard deviation lower than ten quantiles general distribution. Then, the binomial distribution was generated, evaluating the success events and filtering the outliers’ values. For both cases, the filter criterion to select the combination of numerical representation approaches and supervised learning was *success >* 12. This point represents distribution with a probability of success even lower than 0.01, generating significant filter elements. Five combinations of supervised learning algorithms were selected for the case of DNA-binding classification and four for the single-stranded or double-stranded DNA type interaction. The selected methods are summarized in the Table 1. The models present high performances in general, but in all cases, the differences between training and validation metrics allow us to infer that there is overfitting during the training process.

**Table 1:** Selected combinations of supervised learning algorithms and numerical representation approaches for all tasks explored in this work.

| Task | Algorithm | Encoder | Recall |
| --- | --- | --- | --- |
| DNA-binding classification | ExtraTrees | prottrans Uniref | 0.93 |
|  | ExtraTrees | prottrans bdf | 0.93 |
|  | Gradient Boosting | prottrans Uniref | 0.91 |
|  | KNeighbors | prottrans Uniref | 0.93 |
|  | RandomForest | prottrans Uniref | 0.93 |
| Single-stranded or double-stranded | ExtraTrees | prottrans Uniref | 0.90 |
|  | ExtraTrees | prottrans XLU50 | 0.90 |
|  | Gaussian Process | prottrans XLU50 | 0.89 |
|  | Support Vector Classifier | prottrans XLU50 | 0.90 |

All selected combinations of supervised learning algorithms and numerical representation strategies were optimized using the Optuna library (Akiba et al., 2019). Then, two models were selected based on the selected criterion exposed in the pipeline description in the previous section. In the case of DNA-binding, the combination of ExtraTrees supervised learning algorithm and prottrans Uniref was selected. In contrast, the algorithm for the single-stranded or double-stranded identification model was the same, but the pre-trained model selected was prottrans XLU50. The performances obtained by the models are 95% of precision with a Matthews correlation coefficient (MCC) of 0.89 for the DNA-binding models and 89% of precision with an MCC of 0.81 for the single-stranded or double-stranded model.

Figure 3 summarises both constructed models. The confusion matrix (See Figure 3 A and 3 C) describes the capability of the models to identify correctly positive and negative classes. In the case of DNA-binding models, the True Positive and True Negative are higher than the trained model for the single-stranded or double-stranded task, demonstrating a complex process in recognising different types of interactions. Besides, the precision-recall curves were estimated (See Figure 3 B and 3 D) and the average precision was calculated, obtaining a 0.98 for DNA-binding models and a 0.96 for the single-stranded or double-stranded task. This difference is concordant with the confusion matrix, demonstrating the complexity of differentiating between single-stranded and double-stranded type interactions. Finally, a Receiver operating characteristics (ROC) curve was estimated during the training process with *k* = 5 cross-validation for both models (See Figure 3 B and 3 D). Also, the area under curve score was estimated, achieving a 0.98 for the DNA-binding model and a 0.97 for the single-stranded or double-stranded type interaction, demonstrating a high capability of the trained models to recognize DNA-binding proteins and to identify the interaction type.

**Figure 3:**
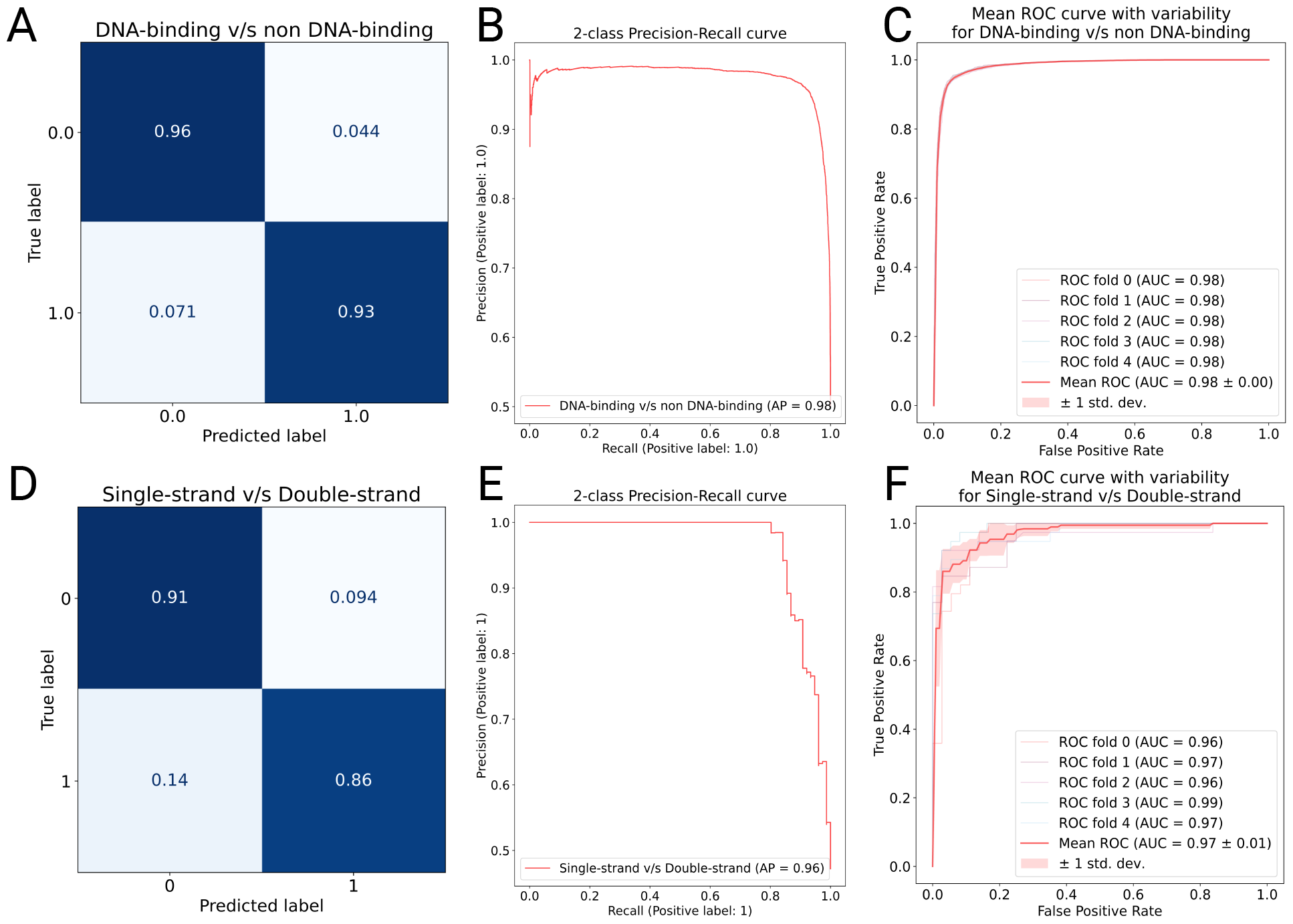
Description through different performances visualization the selected and optimized models for both tasks explored in this work. **A-D** Confusion matrix estimated during the validation process for DNA-binding task single-stranded or double-stranded task, respectively. **B-E** Precision-recall curve estimated during the validation process for DNA-binding task single-stranded or double-stranded task, respectively. The average precision (AP) was calculated in both cases, achieving 0.98 and 0.96, respectively. **C-F** Receiver operating characteristic (ROC) curve estimated during the training process for DNA-binding task single-stranded or double-stranded task, respectively. In both cases, the area under the curve (AUC) was estimated to achieve 0.98 and 0.97, respectively.

A state-of-the-art comparison for both trained models incorporated in RUDEUS is summarised in Table 2. In the case of DNA-binding models, the proposed model achieves the highest specificity values (95.5%) and the highest MCC (0.89) compared to the other methods reported in the literature. The method proposed by (Zhang et al., 2021a) achieves the highest sensibility values. However, the sensitivity value achieved in this work is only 0.1% different from the highest values. In contrast, the single-stranded or double-stranded classification model proposed in this work only achieved the highest MCC value (0.81). Regarding sensitivity, the method proposed by (Ali et al., 2020) has the highest value with 94.2%. Besides, for the specificity values, the method implemented by (Tan et al., 2019) has the highest value with 97.5%. However, both methods show overfitting denoted by the high difference in specificity and sensitivity, respectively, and also denoted by the low MCC value, compared with the MCC obtained in this work.

**Table 2:** State-of-the-art comparison for DNA-binding classification models and single-stranded or doublestranded interaction models.

| Task | Classifier | SN(%) | SP(%) | MCC | Reference |
| --- | --- | --- | --- | --- | --- |
| DNA-binding | RF | 79.3 | 89.0 | 0.69 | (Kumar et al., 2009) |
|  | RF | 83.7 | 90.0 | 0.72 | (Ma et al., 2016) |
|  | SVM | 87.0 | 85.5 | 0.72 | (Zaman et al., 2017) |
|  | SVM | 89.1 | 88.8 | 0.78 | (Ali et al., 2018) |
|  | SVM | 94.1 | 97.6 | 0.92 | (Rahman et al., 2018) |
|  | SVM | 91.1 | 88.8 | 0.79 | (Mishra et al., 2019) |
|  | SVM | 91.8 | 93.0 | 0.84 | (Ali et al., 2019) |
|  | SVM | **93.4** | 93.4 | 0.86 | (Zhang et al., 2021a) |
|  | **ExtraTrees** | 93.3 | **95.5** | **0.89** | **This work** |
| Single-stranded<br>or double-stranded | RF | 90.8 | 78.8 | 0.64 | (Wang et al., 2017) |
|  | SVM | **94.2** | 80.33 | 0.72 | (Ali et al., 2020) |
|  | GTB | 78.4 | **97.5** | 0.79 | (Tan et al., 2019) |
|  | HMM | 85.3 | 92.8 | 0.78 | (Sharma et al., 2021) |
|  | **ExtraTrees** | 87.8 | 91.6 | **0.81** | **This work** |

### 3.3 RUDEUS facilitate the exploration of single-stranded or double-stranded interaction evaluation

More than 20000 protein sequences annotated as DNA-binding proteins were classified as single-stranded or double-stranded using the exploration module available in RUDEUS. First, protein sequences were numerically represented, applying the selected pre-trained models for the strand interaction model. Then, the classification model was applied, and the predictions were obtained. More than 18000 protein sequences were classified as double-stranded interactions. In contrast, only 2000 proteins were identified as single-stranded. The proportions are very similar to those presented in the dataset used to train the predictive model.

Three DNA-binding proteins with an identified strand were evaluated using the bioinformatic structural pipeline previously described in the methodology section. Figure 4 shows the DNA-protein molecular docking obtained, incorporating a general visualization and a specific analysis of the relevant sites for the interaction. The three DNA-binding proteins analyzed in this work have direct interaction mechanisms and have been previously reported in the literature.

**Figure 4:**
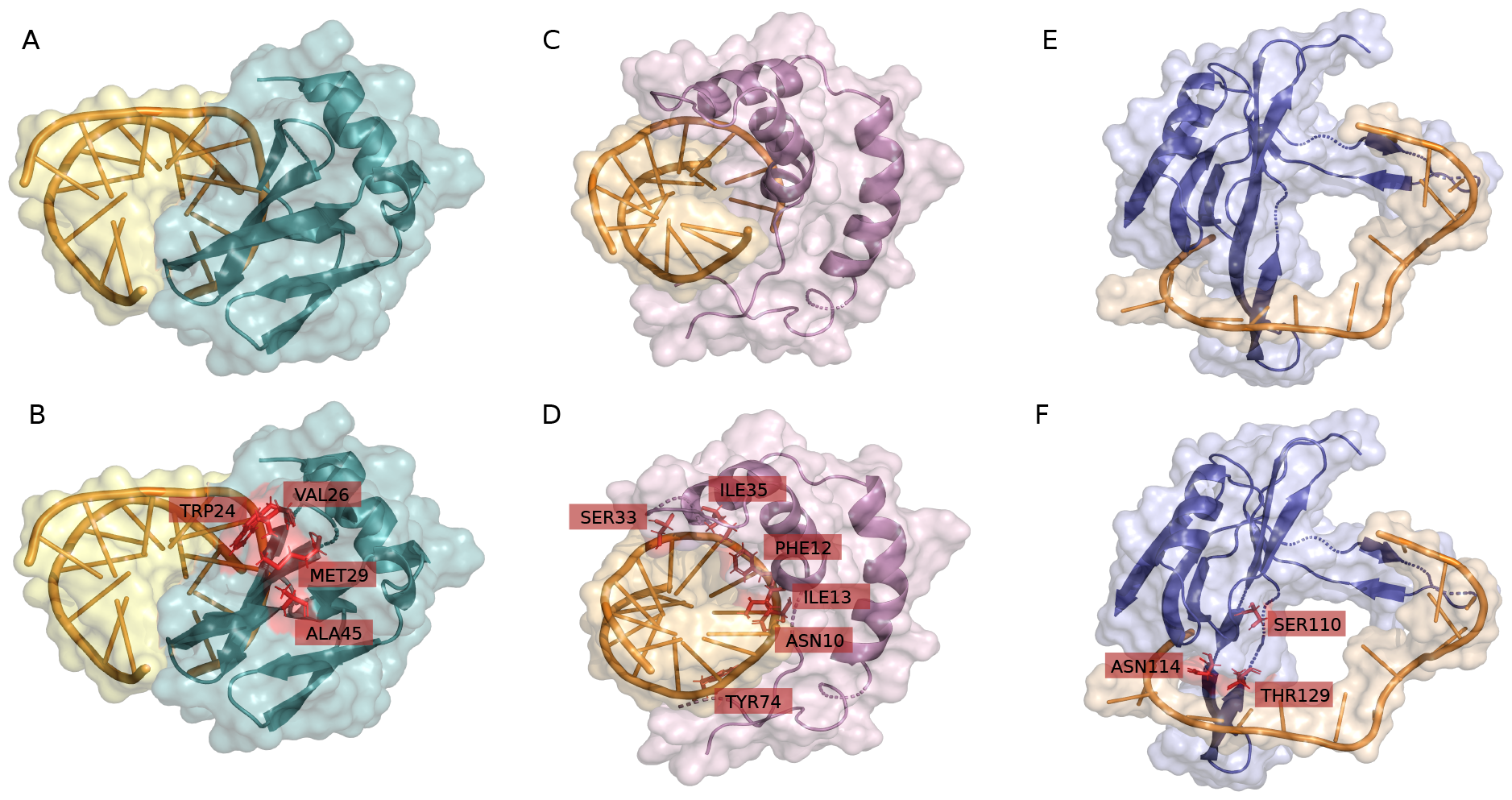
Structural bioinformatics validation through DNA-protein molecular docking for three DNA-binding proteins and their interaction type identified with the models available in RUDEUS. **A-B** DNA-protein molecular docking and the most relevant identified residues for the DNA interaction for the protein 1BNZ. **C-D** DNA-protein molecular docking and the most relevant identified residues for the interaction for the protein 1HRY. **E-F** DNA-protein molecular docking and the most relevant identified residues for the interaction for the protein 3ULP.

Figure 4 A shows the DNA-protein molecular docking for the protein 1BNZ. This hyperthermophile chromosomal protein is reported as a double-stranded union with DNA (Gao et al., 1998; Guagliardi et al., 2002). The hydrophobic residues TRP24, VAL26, MET29, and ALA45 are present on the surface and play an essential role in the DNA binding process (See Figure 4 B). Furthermore, the triple-stranded *β*-sheet (*β*3-*β*4-*β*5) establishes interactions through direct hydrogen bonds/salt bridges, as well as close van der Waals contacts and possible hydrogen bonds/salt bridges in this region, similar to reported in (Gao et al., 1998). Similarly, Figure 4 C shows the DNA-protein molecular docking for the protein 1HRY. This protein has the function of terminating the male sex by being a transitional activator of the gene responsible for the regression of the female Müllerian ducts in male embryos and modulating the genetic activity associated with sexual differentiation (Werner et al., 1995). 1HRY has six residues (ASN10, PHE12, ILE13, SER33, ILE35, SER36, and TYR74) (See Figure 4 D) that come into contact with the DNA bases. In the case of SER33, the carboxyl group bonds with G9, while the carboxyamide group of ASN10 is involved in electrostatic interactions with G13, C4 and A5. Additionally, the hydroxyl group of TYR74 can form a hydrogen bond with T14, as previously described in (Werner et al., 1995).

In contrast, Figure 4 E shows the DNA-protein molecular docking for the protein 3ULP. This protein is called Pf-SSB. This single-strand DNA-binding protein plays an essential role in the DNA metabolism of the eukaryotic parasite responsible for malaria. The single-stranded DNA binding protein (SSB) is encoded in the nucleus and transported to the apicoplast, where it is part of the replication and maintenance of the apicoplast genome (Antony et al., 2012). The 3ULP protein exhibits a stable structure in the form of a homotetramer. Despite this, the residues that come into contact with the DNA are identical in all four subunits, each composed of an N-terminal OB-fold domain, which contains the binding site, and an unstructured C-terminal (Antony et al., 2012). In particular, residues S110, N114 and T129 contact the DNA curvature (See Figure 4 F).

## 4 Conclusion

This work presents RUDEUS, a Python library designed to investigate DNA-binding proteins, to identify this function and assess DNA-strand recognition types. The methodology involves a versatile pipeline incorporating protein language models, supervised learning algorithms, and a Bayesian optimization strategy to train classification models. These models exhibit superior performance, outperforming state-of-the-art benchmarks in sensitivity, specificity, and MCC scores. Despite this success, the simplicity and replicability of the previously established method remain advantageous.

An exploration process was executed to showcase RUDEUS’s utility, enabling the annotation of over 20,000 protein sequences as single-stranded or double-stranded through structural bioinformatic approaches, validated by DNA-protein molecular docking. RUDEUS’s user-friendly interface and robust functionality enhance its applicability, facilitating integration into protein design pipelines such as landscape reconstruction, directed evolution, and latent space exploration via deep generative models.

## 5 Author contributions statement

IM-B and DM-O: conceptualization. DM-O, GC-M, and NS-G: methodology. DM-O and RU-P: validation. IM-B, GC-M, and NS-G: investigation. DM-O, IM-B, RU-P, and GC-M: writing, review, and editing. DM-O and RU-P: supervision and funding resources. DM-O: project administration.

## Acknowledgments

This research has been financed mainly by the Centre for Biotechnology and Bioengineering - CeBiB (PIA project FB0001, Conicyt, Chile). DM-O acknowledges ANID for the project “SUBVENCIÓN A INSTALACIÓN EN LA ACADEMIA CONVOCATORIA A Ñ O 2022”, Folio 85220004. RU-P acknowledges ANID for the grant Fondecyt 1230298.

## Codes and data availability statement

RUDEUS is available for non-commercial use under the MIT License. The source code, models, and implemented strategies are all available in the GitHub repository https://github.com/ProteinEngineering-PESB2/RUDEUS.

## References

Akiba, T., Sano, S., Yanase, T., Ohta, T., and Koyama, M. (2019). Optuna: A next-generation hyperparameter optimization framework. In Proceedings of the 25th ACM SIGKDD international conference on knowledge discovery & data mining, pages 2623–2631.

Alendar, A. and Berns, A. (2021a). Sentinels of chromatin: chromodomain helicase dna-binding proteins in development and disease. Genes & Development, 35(21-22):1403–1430.

Alendar, A. and Berns, A. (2021b). Sentinels of chromatin: chromodomain helicase dna-binding proteins in development and disease. Genes & Development, 35(21-22):1403–1430.

Ali, F., Ahmed, S., Swati, Z. N. K., and Akbar, S. (2019). Dp-binder: machine learning model for prediction of dna-binding proteins by fusing evolutionary and physicochemical information. Journal of Computer-Aided Molecular Design, 33:645–658.

Ali, F., Arif, M., Khan, Z. U., Kabir, M., Ahmed, S., and Yu, D.-J. (2020). Sdbp-pred: Prediction of single-stranded and double-stranded dna-binding proteins by extending consensus sequence and k-segmentation strategies into pssm. Analytical biochemistry, 589:113494.

Ali, F., Kabir, M., Arif, M., Swati, Z. N. K., Khan, Z. U., Ullah, M., and Yu, D.-J. (2018). Dbppred-pdsd: Machine learning approach for prediction of dna-binding proteins using discrete wavelet transform and optimized integrated features space. Chemometrics and Intelligent Laboratory Systems, 182:21–30.

Ali, F., Kumar, H., Patil, S., Ahmed, A., Banjar, A., and Daud, A. (2022). Dbp-deepcnn: prediction of dna-binding proteins using wavelet-based denoising and deep learning. Chemometrics and Intelligent Laboratory Systems, 229:104639.

Antony, E., Weiland, E. A., Korolev, S., and Lohman, T. M. (2012). Plasmodium falciparum ssb tetramer wraps single-stranded dna with similar topology but opposite polarity to e. coli ssb. Journal of molecular biology, 420(4-5):269–283.

Arora, S., Gupta, S., Verma, S., and Malik, I. (2023). Prediction of dna interacting residues. In 2023 International Conference on Computational Intelligence, Communication Technology and Networking (CICTN), pages 54–57. IEEE.

Attali, I., Botchan, M. R., and Berger, J. M. (2021). Structural mechanisms for replicating dna in eukaryotes. Annual review of biochemistry, 90:77–106.

Banjar, A., Ali, F., Alghushairy, O., and Daud, A. (2022). idbp-pbmd: A machine learning model for detection of dna-binding proteins by extending compression techniques into evolutionary profile. Chemometrics and Intelligent Laboratory Systems, 231:104697.

Barukab, O., Ali, F., Alghamdi, W., Bassam, Y., and Khan, S. A. (2022). Dbp-cnn: Deep learning-based prediction of dna-binding proteins by coupling discrete cosine transform with two-dimensional convolutional neural network. Expert Systems with Applications, 197:116729.

Dallago, C., Schütze, K., Heinzinger, M., Olenyi, T., Littmann, M., Lu, A. X., Yang, K. K., Min, S., Yoon, S., Morton, J. T., and Rost, B. (2021). Learned embeddings from deep learning to visualize and predict protein sets. Current Protocols, 1(5):e113.

Elnaggar, A., Heinzinger, M., Dallago, C., Rihawi, G., Wang, Y., Jones, L., Gibbs, T., Feher, T., Angerer, C., Steinegger, M., Bhowmik, D., and Rost, B. (2020). Prottrans: Towards cracking the language of life’s code through self-supervised deep learning and high performance computing.

Fernández, D., Olivera-Nappa, Á., Uribe-Paredes, R., and Medina-Ortiz, D. (2023). Exploring machine learning algorithms and protein language models strategies to develop enzyme classification systems. In International Work-Conference on Bioinformatics and Biomedical Engineering, pages 307–319. Springer.

Fu, Y., Ling, Z., Arabnia, H., and Deng, Y. (2020). Current trend and development in bioinformatics research.

Gao, Y.-G., Su, S.-Y., Robinson, H., Padmanabhan, S., Lim, L., McCrary, B. S., Edmondson, S. P., Shriver, J. W., and Wang, A. H.-J. (1998). The crystal structure of the hyperthermophile chromosomal protein sso7d bound to dna. Nature structural biology, 5(9):782–786.

Guagliardi, A., Cerchia, L., Rossi, M., et al. (2002). The sso7d protein of sulfolobus solfataricus: in vitro relationship among different activities. Archaea, 1:87–93.

Gupta, N. K., Wilkinson, E. A., Karuppannan, S. K., Bailey, L., Vilan, A., Zhang, Z., Qi, D.-C., Tadich, A., Tuite, E. M., Pike, A. R., et al. (2021). Role of order in the mechanism of charge transport across single-stranded and double-stranded dna monolayers in tunnel junctions. Journal of the American Chemical Society, 143(48):20309–20319.

Hu, S., Ma, R., and Wang, H. (2019). An improved deep learning method for predicting dna-binding proteins based on contextual features in amino acid sequences. PLoS one, 14(11):e0225317.

Kabir, A., Bhattarai, M., Rasmussen, K. O., Shehu, A., Bishop, A. R., Alexandrov, B. S., and Usheva, A. (2024). Advancing transcription factor binding site prediction using dna breathing dynamics and sequence transformers via cross attention. bioRxiv, pages 2024–01.

Kumar, K. K., Pugalenthi, G., and Suganthan, P. N. (2009). Dna-prot: identification of dna binding proteins from protein sequence information using random forest. Journal of Biomolecular Structure and Dynamics, 26(6):679–686.

Lye, Y. S. and Chen, Y.-R. (2022). Tar dna-binding protein 43 oligomers in physiology and pathology. IUBMB life, 74(8):794–811.

Ma, X., Guo, J., and Sun, X. (2016). Dnabp: Identification of dna-binding proteins based on feature selection using a random forest and predicting binding residues. PloS one, 11(12):e0167345.

McKinney, W. et al. (2011). pandas: a foundational python library for data analysis and statistics. Python for high performance and scientific computing, 14(9):1–9.

Medina, D., Sepulveda-Yanez, J., Alvarez-Saravia, D., Uribe-Paredes, R., Veelken, H., and Navarrete, M. (2023). Artificial intelligence approach for the discovery of autoantigen recognition by b-cell lymphomas. Blood, 142:125.

Medina-Ortiz, D., Contreras, S., Amado-Hinojosa, J., Torres-Almonacid, J., Asenjo, J. A., Navarrete, M., and Olivera-Nappa, Á. (2022). Generalized property-based encoders and digital signal processing facilitate predictive tasks in protein engineering. Frontiers in Molecular Biosciences, 9.

Medina-Ortiz, D., Contreras, S., Quiroz, C., and Olivera-Nappa, Á. (2020). Development of supervised learning predictive models for highly non-linear biological, biomedical, and general datasets. Frontiers in molecular biosciences, 7:13.

Meier, J., Rao, R., Verkuil, R., Liu, J., Sercu, T., and Rives, A. (2021). Language models enable zero-shot prediction of the effects of mutations on protein function. In Ranzato, M., Beygelzimer, A., Dauphin, Y., Liang, P., and Vaughan, J. W., editors, Advances in Neural Information Processing Systems, volume 34, pages 29287–29303. Curran Associates, Inc.

Mishra, A., Pokhrel, P., and Hoque, M. T. (2019). Stackdppred: a stacking based prediction of dna-binding protein from sequence. Bioinformatics, 35(3):433–441.

Pan, Y., Zhou, S., and Guan, J. (2020). Computationally identifying hot spots in protein-dna binding interfaces using an ensemble approach. BMC bioinformatics, 21:1–16.

Pedregosa, F., Varoquaux, G., Gramfort, A., Michel, V., Thirion, B., Grisel, O., Blondel, M., Prettenhofer, P., Weiss, R., Dubourg, V., et al. (2011). Scikit-learn: Machine learning in python. the Journal of machine Learning research, 12:2825–2830.

Rahman, M. S., Shatabda, S., Saha, S., Kaykobad, M., and Rahman, M. S. (2018). Dpp-pseaac: a dna-binding protein prediction model using chou’s general pseaac. Journal of theoretical biology, 452:22–34.

Rives, A., Meier, J., Sercu, T., Goyal, S., Lin, Z., Liu, J., Guo, D., Ott, M., Zitnick, C. L., Ma, J., and Fergus, R. (2021). Biological structure and function emerge from scaling unsupervised learning to 250 million protein sequences. Proceedings of the National Academy of Sciences, 118(15).

Roel-Touris, J., Bonvin, A. M., and Jim énez-García, B. (2020). Lightdock goes information-driven. Bioinformatics, 36(3):950–952.

Shadab, S., Khan, M. T. A., Neezi, N. A., Adilina, S., and Shatabda, S. (2020). Deepdbp: deep neural networks for identification of dna-binding proteins. Informatics in Medicine Unlocked, 19:100318.

Sharma, R., Kumar, S., Tsunoda, T., Kumarevel, T., and Sharma, A. (2021). Single-stranded and double-stranded dna-binding protein prediction using hmm profiles. Analytical biochemistry, 612:113954.

Tan, C., Wang, T., Yang, W., and Deng, L. (2019). Predpsd: a gradient tree boosting approach for single-stranded and double-stranded dna binding protein prediction. Molecules, 25(1):98.

Wang, W., Sun, L., Zhang, S., Zhang, H., Shi, J., Xu, T., and Li, K. (2017). Analysis and prediction of single-stranded and double-stranded dna binding proteins based on protein sequences. BMC bioinformatics, 18:1–10.

Wang, W., Zhang, Y., Liu, D., Zhang, H., Wang, X., and Zhou, Y. (2022a). Prediction of dna-binding protein–drug-binding sites using residue interaction networks and sequence feature. Frontiers in Bioengineering and Biotechnology, 10:822392.

Wang, Y., Zhang, L., Huang, T., Wu, G.-R., Zhou, Q., Wang, F.-X., Chen, L.-M., Sun, F., Lv, Y., Xiong, F., et al. (2022b). The methyl-cpg-binding domain 2 facilitates pulmonary fibrosis by orchestrating fibroblast to myofibroblast differentiation. European Respiratory Journal, 60(3).

Wang, Z., Gong, M., Liu, Y., Xiong, S., Wang, M., Zhou, J., and Zhang, Y. (2022c). Towards a better understanding of tf-dna binding prediction from genomic features. Computers in Biology and Medicine, 149:105993.

Werner, M. H., Huth, J. R., Gronenborn, A. M., and Clore, G. M. (1995). Molecular basis of human 46x, y sex reversal revealed from the three-dimensional solution structure of the human sry-dna complex. Cell, 81(5):705–714.

Zaman, R., Chowdhury, S. Y., Rashid, M. A., Sharma, A., Dehzangi, A., Shatabda, S., et al. (2017). Hmmbinder: Dna-binding protein prediction using hmm profile based features. BioMed research international, 2017.

Zhang, J., Chen, Q., and Liu, B. (2020). idrbp mmc: identifying dna-binding proteins and rna-binding proteins based on multi-label learning model and motif-based convolutional neural network. Journal of molecular biology, 432(22):5860–5875.

Zhang, Q., Liu, P., Wang, X., Zhang, Y., Han, Y., and Yu, B. (2021a). Stackpdb: predicting dna-binding proteins based on xgb-rfe feature optimization and stacked ensemble classifier. Applied Soft Computing, 99:106921.

Zhang, Y., Bao, W., Cao, Y., Cong, H., Chen, B., and Chen, Y. (2022). A survey on protein–dna-binding sites in computational biology. Briefings in Functional Genomics, 21(5):357–375.

Zhang, Y., Wang, Z., Zeng, Y., Zhou, J., and Zou, Q. (2021b). High-resolution transcription factor binding sites prediction improved performance and interpretability by deep learning method. Briefings in Bioinformatics, 22(6):bbab273.

